# Cellular variability of nonsense-mediated mRNA decay

**DOI:** 10.1101/2021.03.31.437867

**Authors:** Hanae Sato, Robert H. Singer

## Abstract

Nonsense-mediated mRNA decay (NMD) is an mRNA degradation pathway that eliminates transcripts containing premature termination codons (PTCs). Half-lives of the mRNAs containing PTCs demonstrate that a small percent escape surveillance and do not degrade. It is not known whether this escape represents variable mRNA degradation within cells or, alternatively cells within the population are resistant. Here we demonstrate a single-cell approach with a bi-directional reporter, which expresses two β-globin genes with or without a PTC in the same cell, to characterize the efficiency of NMD in individual cells. We found a broad range of NMD efficiency in the population; some cells degraded essentially all of the mRNAs, while others escaped NMD almost completely. Characterization of NMD efficiency together with NMD regulators in single cells showed cell-to-cell variability of NMD reflects the differential level of surveillance factors, SMG1 and phosphorylated UPF1. A single-cell fluorescent reporter system that enabled detection of NMD using flow cytometry revealed that this escape occurred either by translational readthrough at the PTC or by a failure of mRNA degradation after successful translation termination at the PTC.

## Introduction

Nonsense-mediated mRNA decay (NMD) is a highly conserved mRNA degradation pathway in eukaryotes. NMD eliminates aberrant mRNAs containing premature termination codons (PTC) that potentially produce truncated dominant-negative or gain-of-function proteins (Holbrook et al., 2004) as well as transcripts containing introns in their 3’ untranslated region (UTR) or longer 3’UTRs (Colombo et al., 2017; Kervestin and Jacobson, 2012; Mendell et al., 2004). A major research focus has been to understand how NMD recognizes its target transcripts. In mammalian cells, two models have been suggested to explain selective mRNA degradation in NMD: the “EJC-model”, whereby the existence of splicing-generated exon-junction complexes (EJCs) more than 50 to 55 nucleotides downstream of the PTC define an NMD substrate(Nagy and Maquat, 1998). An alternative model, “faux 3’UTR”, proposes that the distance from the termination codon to the poly (A) tail provides for the targeting of NMD due to the competitive binding of the major NMD regulator, up-frameshift factor 1 (UPF1) and cytoplasmic poly (A) binding protein 1 (PABC1) with eukaryotic polypeptide chain release factor 3 (eRF3), which interacts with eRF1 and recognizes termination codons including the PTC(Amrani et al., 2004). Recent genome-wide exome and transcriptome approaches, which investigated genetic variants predicted to be targeted by NMD and their transcriptome profiles, discovered that a significant proportion of transcripts predicted as NMD targets supports the EJC model (Linde et al., 2007; Rivas et al., 2015).

NMD efficiency varies across transcripts (Gudikote and Wilkinson, 2002) depending on the position of the PTC (Neu-Yilik et al., 2011; Romao et al., 2000; Zhang and Maquat, 1997), tissue type (Linde et al., 2007; Rivas et al., 2015; Zetoune et al., 2008), oncogenesis (Wang et al., 2017) and stress conditions (Gardner, 2008; Karam et al., 2015; Lykke-Andersen and Jensen, 2015; Ottens and Gehring, 2016; Wang et al., 2011; Wengrod et al., 2013). Genome-wide studies also reveal that a significant proportion of transcripts predicted to trigger NMD escape from degradation by unknown mechanisms (Lappalainen et al., 2013) (Rivas et al., 2015) (Lindeboom et al., 2016). Conditional NMD inhibition, “NMD escape”, seems to be a general feature of this process since a subpopulation of NMD substrates escapes from NMD (~30%) (Belgrader et al., 1994; Cheng and Maquat, 1993). However, little is known about how and what leads to NMD escape.

Cell variation is a fundamental characteristic in the process of differentiation, oncogenesis, aging, and many other biological processes (Tanay and Regev, 2017). The technological advances of single-cell approaches such as fluorescence-activated cell sorting (FACS), single-cell RNA sequencing and single molecule FISH have uncovered dramatic variability and diversity in cell populations where heterogeneity was previously unappreciated (Review (Neu et al., 2017; Philipp Angerer, 2017)). The ordinary approaches to determine NMD efficiency, characterizing half-lives of PTC-containing transcripts compared with normal transcripts are generally ensemble assays. These determine the mean of NMD efficiency but are unable to investigate the heterogeneities in the population with single-cell resolution.

## Results

In this study, we established an assay to assess NMD efficiency in individual cells. We used a bidirectional system which allowed the simultaneous and calibrated expression of two transcripts from two open reading frames (ORFs) under the same promoter within the same cell (Trcek et al., 2013). This allowed us to compare genes with or without PTC in the same cell, eliminating extrinsic variability. We confirmed that the PonA bidirectional promoter expressed equally in both directions. We used a construct which expressed wild-type β-globin (Gl) ORF in either direction but contained different stem-loops in the 3’UTR (MS2 or PP7 stem-loops, **Figure 1A&B; WW**), as well as swapping their locations as a control (**Fig.S1; Switch**). These were transiently transfected in human osteosarcoma U2OS PonA cells. After induction for 24-hours, cells were fixed for singlemolecule fluorescence in situ hybridization (smFISH). The individual mRNA molecules were hybridized using fluorescent dye conjugated probes to their respective stem-loops (**Figure 1C & S1**) for fluorescence microscopy. Individual mRNAs were counted using FISH-quant (Mueller et al., 2013), and correlation of expression from either side of the promoter was determined in a scatter plot (**Figure 1E & S1**). The linear correlation between the expression of the mRNAs showed an equivalent induction of transcription from either orientation of the PonA bi-directional promoter. We also found similar results after a one-hour induction (**Figure S2**).

**Figure 1.**
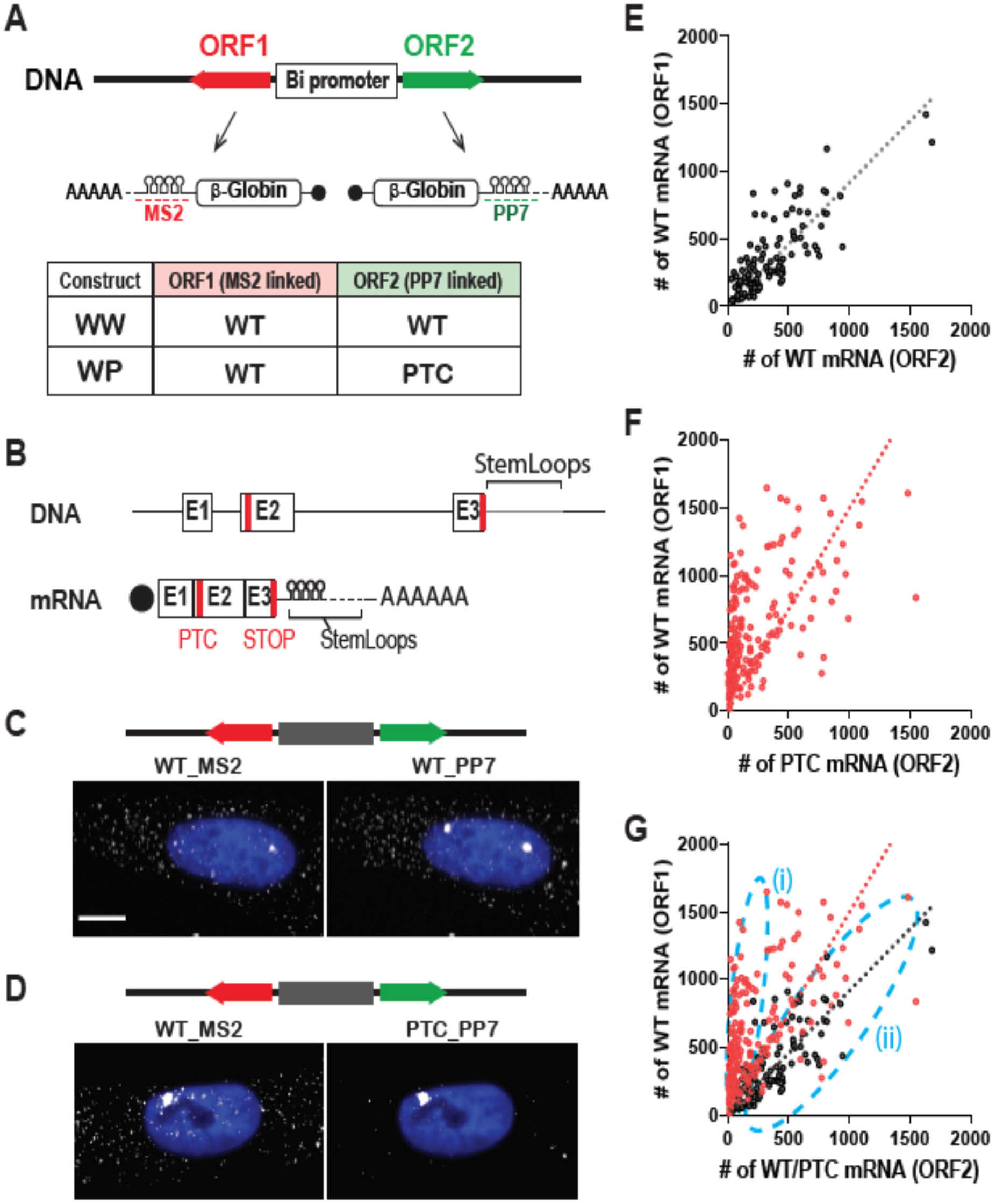
Cellular variability of NMD. (A) Schematic of PonA inducible bi-directional promoter expressing NMD reporter Gl genes. These transcripts contain MS2 or PP7 sequences in the 3’UTR that hybridize with Quasar 670-labeled (red dotted line) or Quasar 570-labeled (green dotted line) FISH probes. The table shows the mRNAs expressing from the bi-directional promoter from each construct. (B) Structures of NMD reporter Gl (DNA; upper and mRNA; lower) containing a PTC at position 39 (PTC). Horizontal lines represent introns, 5’-UTR and 3’-UTR, and boxes represent each of the three Gl exons (E1-3) joined by splicing-generated exon-exon junctions. Black dot, cap structure; Red lines, termination codons; STOP, normal termination codon; AAAAAA, poly(A) tail. (C-D) smFISH images of U2OS PonA cells co-expressing Gl WT (left; expressed from ORF1) and Gl WT (C) or PTC (D) mRNA (expressed from ORF2) from a bi-directional promoter. NMD reporter was transiently expressed. MS2_mRNAs and PP7_mRNAs were simultaneously labeled with Quasar 670-- and Quasar 570-conjugated FISH probes. Note the lack of cytoplasmic mRNA due to the PTC in (D). The nuclei were stained by DAPI (blue). Bar=10um. (E-G) The Scatter plots show the number of Gl WT (ORF1) and Gl WT or PTC (ORF2) mRNAs expressing from WW (E, black), WT (F, red), and both were superimposed in (G). Each dot in scatter plots denotes the number of Gl WT (ORF1; y-axis) and WT or PTC (ORF2; x-axis) mRNAs in a single cell. Dotted lines indicate regression lines. Subpopulations of cells with differential NMD efficiency ((i) efficient NMD and (ii) escape from NMD) are shown as light dotted blue lines.

The approach was then applied to an NMD target combined with a wild type sequence We performed the same experiment with a PTC-containing ORF in one of the orientations (**Figure 1A&D; WP**). Individual mRNAs were counted as described below, then each number of transcripts was represented in a scatter plot (**Figure 1F)**. In contrast to WW transfected cells, we found that WP transfected cells showed a different regression curve with a steeper slope (slope=1.5, R-squared =0.33, **Figure 1F**). This is due to the reduction of PTC containing mRNA by NMD. Notably, the low R-squared value of the regression indicates variable NMD efficiencies, where the population of cells escaping NMD is apparent. (**Figure 1G, (i) and (ii)**). This result demonstrated the variability of NMD efficiency, not detectable using ensemble measurements such as RT-qPCR and northern blotting. The ability to quantify the escaping population allows an analysis of its characteristics.

Since translation is required for NMD, one mechanism to explain how some cells escape from NMD could be from reduction of translational activity. Certain cellular stress conditions such as cellular hypoxia and amino acid deprivation cause translation shut-off through elF2α phosphorylation and result in inactivation of NMD (Gardner, 2008; Karam et al., 2015; Wang et al., 2011; Wengrod et al., 2013). To test if these cells escaped from NMD due to the inactivation of translation, we performed simultaneous detection of NMD efficiency and translational activity using smFISH and the click-it HPG (L-Homopropargylglycine) system. HPG is the amino acid analog of methionine, and the HPG translation detection system is based on the fluorescence labelling by HPG incorporation into nascent proteins. As shown in **Figure 2**, treatment by cycloheximide (CHX), an inhibitor of translation elongation, decreased the detection of HPG incorporation (**Figure S3A&B**), confirming successful reduction of translation activity using this system. The simultaneous detection of NMD efficiency and translation activity showed no correlation (**Figure S3C**). This indicated that cells could escape from NMD even when they are actively translating. NMD efficiency had no correlation with cell size either, indicating that the cell cycle phase, as well as the nuclear size or the level of expression were not involved in the escape (**Figure S4**).

**Figure 2.**
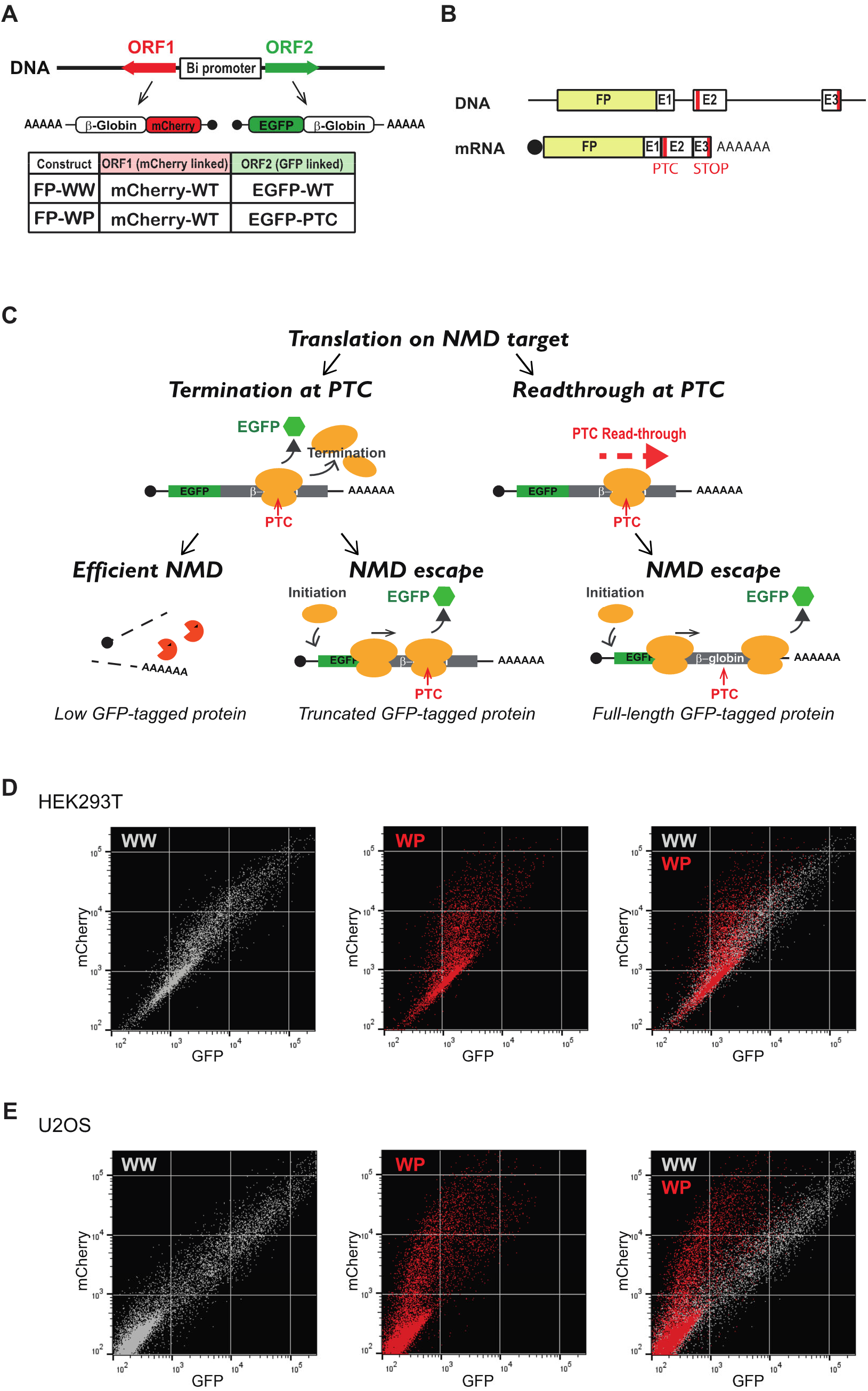
Single-cell analysis of NMD efficiency using flow cytometry. Schematic of translational reporter Gl genes. These transcripts contain EGFP or mCherry coding sequences upstream of the Gl gene. The table shows the mRNAs expressing from the bi-directional promoter from each construct. (B) Structures of Gl DNA (upper) and mRNA (lower) containing a PTC at position 39 (PTC). Horizontal lines represent introns, 5’-UTR and 3’-UTR, and boxes represent each of the three Gl exons (E1-3) joined by splicing-generated exon-exon junctions. Black dot, cap structure; Red lines, termination codons; STOP, normal termination codon; AAAAAA, poly(A) tail; FP, fluorescent protein. (C) Schematic of EGFP expression from NMD reporter construct and expected results. EGFP expresses from the FP_WP construct either when PTC was subject to readthrough or when the PTC was recognized, generating a truncated GFP-containing protein but failed to trigger decay of the mRNA. Low EGFP expression might be detected even when NMD was successfully triggered since EGFP translates prior to PTC recognition. FP construct (WW; Wild-type and Wild-type or WP; Wild-type and PTC) was transiently transfected into (D) HEK293T ponA or (E) U2OS ponA cell line. Transcription was induced with 20nM ponasterone A and the intensities of mCherry and GFP were detected using Area II. 20,000 cells were analyzed and single cells were determined using SCC and FSC, and live cells were selected as DAPI negative cells.

Since PTC recognition is a critical requirement for NMD, translation readthrough might be a key mechanism of NMD escape. Translational readthrough happens due to failure to terminate translation and inserts a near-cognate tRNA for the termination codon that allows elongation to continue. This readthrough at a PTC presumably decreases the likelihood of the mRNA to undergo NMD, and in following rounds of translation of the mRNA since translational readthrough removes the exon junction complexes (EJCs) downstream of the PTC (Sato and Maquat, 2009) required for most of NMD activation (Nagy and Maquat, 1998). NMD escape also may occur due to the failure of the mRNA to degrade after PTC recognition. Therefore, NMD escape may take place in two ways: cells fail to trigger NMD because (1) the ribosome fails to recognize the PTC and reads through it, or (2) translation terminates at a PTC but NMD regulators fail to trigger decay.

To test these two models, we developed a fluorescent NMD reporter system, by fusing EGFP or mCherry coding sequences upstream of either β-globin gene (**Figure 2A-C**). The efficiency of NMD can be monitored as a ratio of the two fluorescence intensities in single cells. This allows the isolation of the specific cell population that exhibits NMD escape by fluorescence-activated cell sorting (FACS). Cells expressing mCherry-tagged wildtype β-globin and EGFP-tagged wild-type or EGFP-tagged PTC-containing β-globin constructs were transiently transfected into HEK293T ponA cells or U2OS ponA cells, and transcription induced by ponasterone A for 24-hours. The intensities of mCherry and EGFP in the cells were determined using flow cytometric analysis. The intensity of mCherry represents the control mRNA; NMD efficiency can be quantified by a ratio of two fluorescence intensities expressed in each cell. Most wild-type cells exhibited both EGFP and mCherry doublepositive and a scatter plot of the fluorescent intensities represented a linear correlation, confirming the equivalent expression of both proteins when expressed from the bidirectional promoter (**Figure 2 D&E middle**). In contrast, the scatter plot of the fluorescent intensities is skewed toward lower EGFP intensities when one of the wild-type mRNAs (EGFP-tagged β-globin) was replaced with the NMD substrate, indicating that these mRNAs were degraded. However, some of the cells demonstrated an EGFP/mCherry ratio more characteristic of wild-type which represented the cells undergoing NMD escape (**Figure 2 D&E middle**). A similar pattern was observed in both HEK293T and U2OS cell lines, suggesting NMD escape in this reporter is common to these cell lines.

NMD involves the core regulator up-frameshift proteins (UPFs). UPF1 initiates the NMD process by interacting with eRF3 at the PTC (Kashima et al., 2006). The interaction of PTC-bound UPF1 with EJCs, which consists of UPF2, UPF3A, or UPF3B, detects the PTC in mammals (Gehring et al., 2005; Gehring et al., 2003; Lejeune et al., 2002). Besides UPFs, NMD involves SMG proteins such as SMG1, SMG5, SMG7, and SMG6 in mammalian cells. The phosphatidylinositol-kinase related kinase (PIKK) SMG1 phosphorylates UPF1 and activates further degradation of mRNA (Kashima et al., 2006; Yamashita et al., 2001). This degradation is carried out by the endonuclease SMG6 (Eberle et al., 2009; Huntzinger et al., 2008) as well as the heterodimer SMG5-SMG7, which recruits the CCR4-NOT deadenylation complex (Chakrabarti et al., 2014; Jonas et al., 2013; Loh et al., 2013) that leads to decapping, deadenylation and exonucleolytic degradation (Nicholson et al., 2018; Unterholzner and Izaurralde, 2004). Since NMD escape could result from reduced expression of one of these known NMD regulators in individual cells, we correlated NMD efficiency with protein levels of NMD regulators, UPF1, SMG1, and SMG6, along with the phosphorylation of UPF1, using our imaging approach. The relevant construct was transiently transfected into U2OS ponA cells and induced for 24-hours. UPF1, UPF1 phosphorylation, SMG1, or SMG6 were labeled by immunofluorescence using their specific antibodies followed by anti-rabbit Alexa Fluor 647. The fluorescence levels of EGFP, mCherry and IF labeling NMD regulators were detected by fluorescence microscopy and their intensities in each cell were determined using CellProfiler (Carpenter et al., 2006; Kamentsky et al., 2011; McQuin et al., 2018). The results showed that NMD efficiency was correlated with the level of SMG1 (**Figure 3A&B**); less correlation was found with the level of UPF1 or with SMG6 (**Figure S5**). A correlation of NMD efficiency (a negative correlation with NMD escape) with phosphorylation of UPF1 was also evident (**Figure 3C**). This is likely due to the reduction of SMG1 which phosphorylates UPF1. The negative correlation of NMD efficiency with the level of SMG1 is consistent with previous work showing the overexpression or downregulation of SMG1 increases or decreases NMD efficiency respectively (Usuki et al., 2006; Yamashita et al., 2001).

**Figure 3.**
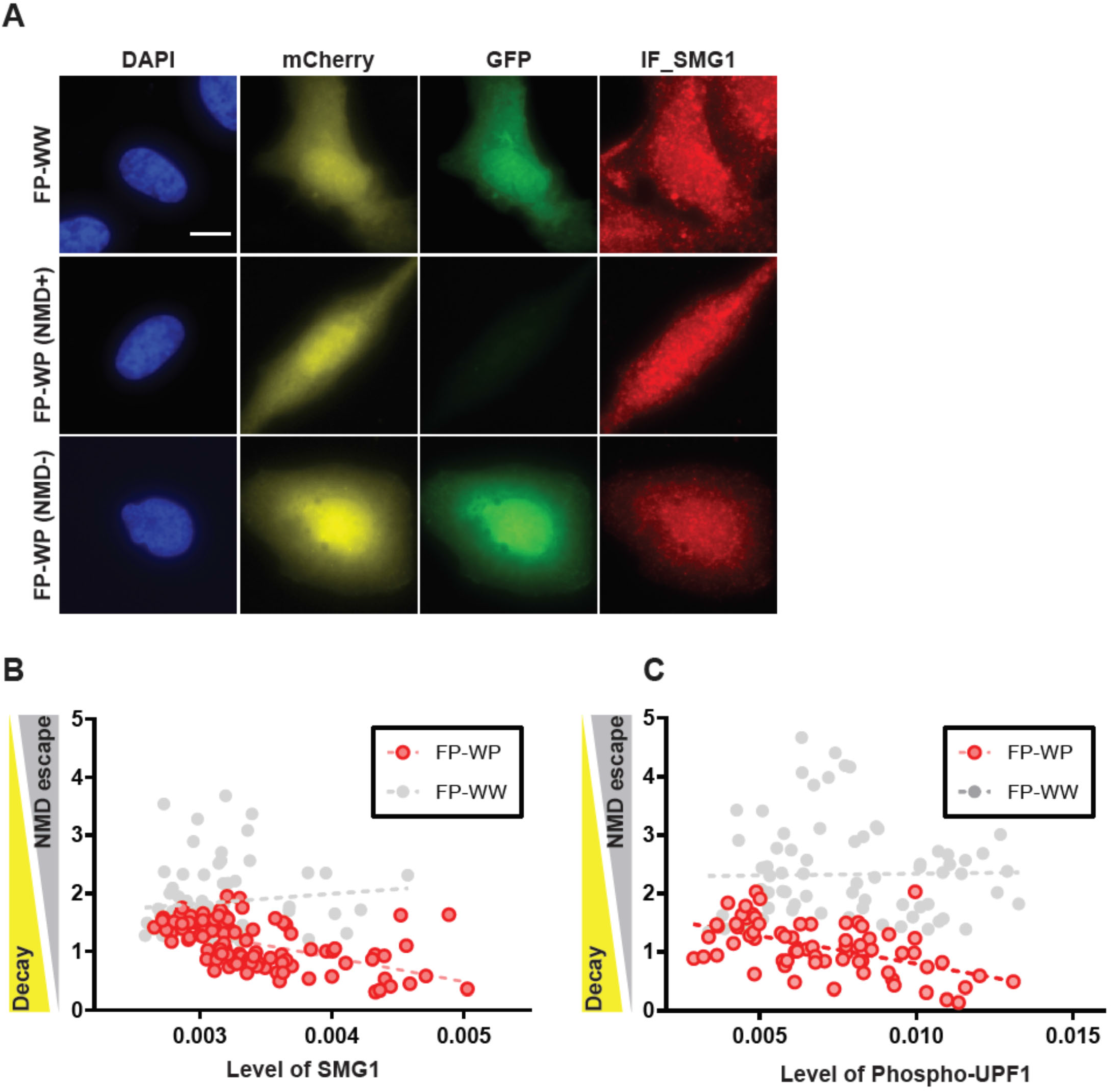
Correlation of NMD efficiency with NMD regulator SMG1. (A) Simultaneous detection of Fluorescence NMD reporter and NMD regulator, SMG1. FP construct (FP-WW; Wild-type and Wild-type or FP-WP; Wild-type and PTC) was transiently transfected into U2OS ponA cell line. Transcription was induced with 20nM ponasterone A for 24 hours. The level of SMG1 was determined by immunofluorescence using anti-SMG1 antibody followed by Alexa Fluor 647 linked anti-rabbit IgG. The intensity of mCherry, EGFP, and the level of SMG1 (IF_SMG1), DAPI-stained nuclei were detected using fluorescence microscopy. Bar = 10 μm. (B) Correlation of NMD efficiency and the level of SMG1. Pearson r=-0.5597, P (twotailed) <0.0001, n=116. (C) Correlation of NMD efficiency and the level of phosphorylation of UPF1. Pearson r=-0.5347, P (two-tailed) <0.0001, n=68. (B&C) Mean intensity of EGFP, mCherry and IF representing the level of SMG1 or phosphorylation of UPF1 in individual cells were determined using Cellprofiler software (Carpenter et al., 2006; Kamentsky et al., 2011; McQuin et al., 2018). The NMD efficiency in single cells was calculated by the intensity of EGFP normalized by the intensity of mCherry (EGFP/mCherry ratio). Single dots denote fluorescence intensity if IF and normalized NMD efficiency from single cells. X- or y-axis shows mean intensity of IF or NMD efficiency. No correlation between EGFP/mCherry ratio and the level of SMG1 or phosphorylation of UPF1 was found in WW expressing cells. The dotted line indicates the regression line. The correlation test between NMD escape and the level of SMG1 or phosphp-UPF1 was performed by Graphpad prism software.

SMG1 is a multitasking player in the maintenance of telomeres, and the regulation of apoptosis and several cellular stress responses including DNA damage, oxidative and hypoxic stresses (Brown et al., 2011; Brumbaugh et al., 2004; Masse et al., 2008; Nergadze et al., 2004; Oliveira et al., 2008). The interplay between SMG1 and ataxia-telangiectasia mutated (ATM) protein, which is also a PIKK family member, phosphorylates its downstream target p53 on serine 15 and coordinates downstream stress-induced signaling pathways in response to genotoxic stress (Brumbaugh et al., 2004; Gehen et al., 2008; Gewandter et al., 2011). NMD efficiencies may also correlate with ATM autophosphorylation at Ser 1981 (ATM S1981P) which is a common DNA double-strand break (DSB) marker. To test if NMD escape is also correlated with DNA stress response, we employed IF against ATM S1981P using a specific antibody and fluorescence intensities were detected together with an NMD fluorescence reporter in U2OS ponA cells. Interestingly, the imaging approach revealed that subcellular localization of ATM S1981P changed from punctate bright spots (**Figure 4A (a)**) to a homogeneous distribution (**Figure 4A (b)**) in the nucleus when cells escaped from NMD. This uniform distribution pattern of ATM S1981P in the nucleus is consistent with the ATM activation under DSBs (Crescenzi et al., 2008). Additionally, a correlation test between NMD efficiency with the level of ATM in cells did not indicate a correlation (**Figure S6**), however, the level of ATM S1981P did correlate with NMD efficiency and it was translocated to the nucleus in escaped cells (**Figure 4B**). This suggests a link between NMD efficiency and DNA damage stress response.

**Figure 4.**
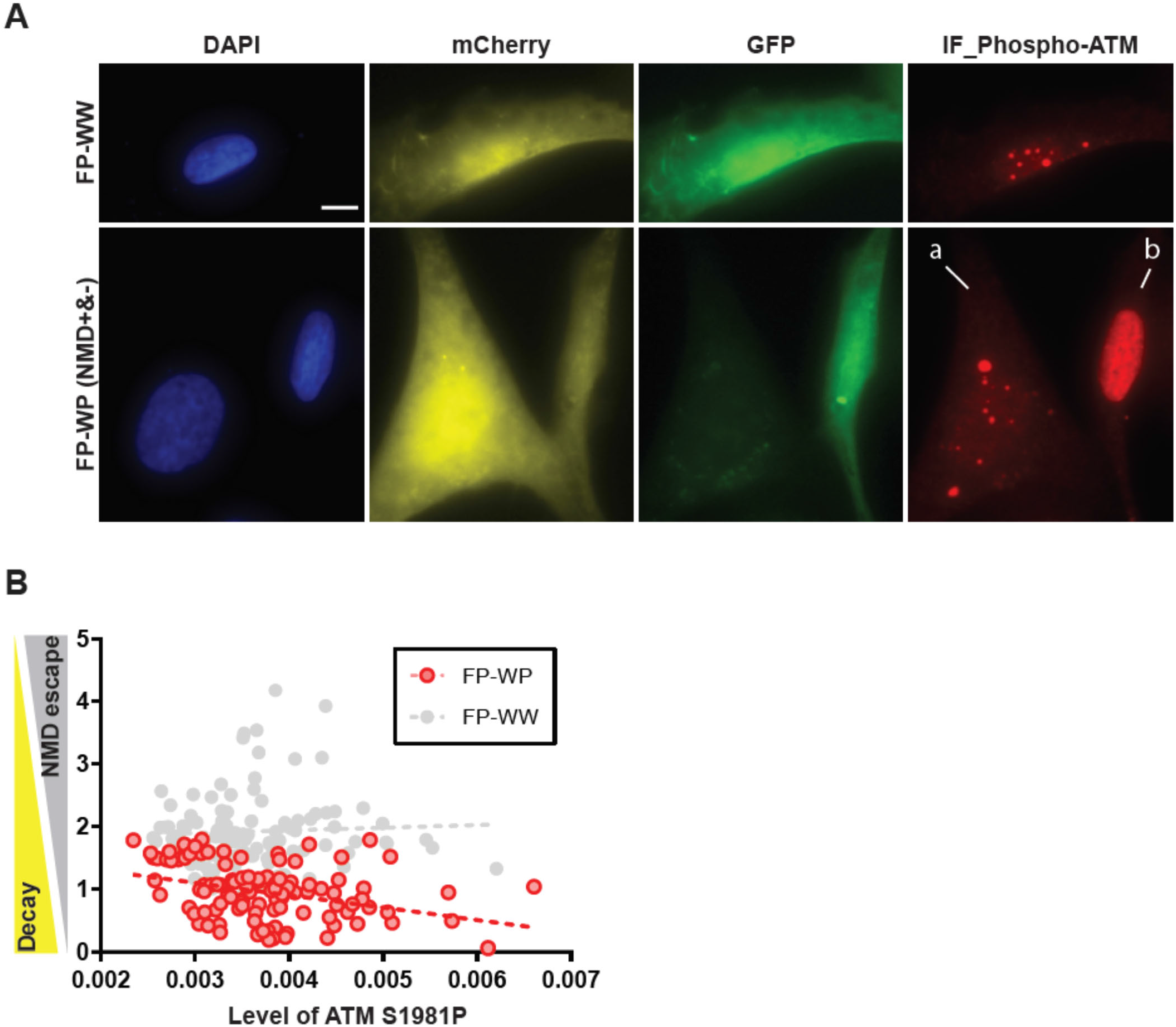
Correlation of NMD efficiency with ATM phosphorylation at S1981. (A) Simultaneous detection of Fluorescence NMD reporter and ATM phosphorylation at S1981. FP construct *(WW;* Wild-type and Wild-type or WP; Wild-type and PTC) was transiently transfected into U2OS ponA cell line. Transcription was induced with 20nM ponasterone A for 24 hours. The level of ATM S1981P was determined by immunofluorescence using anti-ATM S1981P antibody followed by Alexa Fluor 647 linked anti-mouse IgG. The intensity of mCherry, EGFP, DAPI-stained nuclei, or the level of ATM S1981P was detected using fluorescence microscopy. Bar = 10 μm. (a) shows a cell with efficient NMD (lower EGFP expression) and (b) shows a cell with NMD escape (higher EGFP expression). (B) Correlation of NMD efficiency and the level of ATM phosphorylation at S1981. Pearson r=-0.5347, P (two-tailed) <0.0001, n=68. Mean intensity of EGFP, mCherry and IF representing the level of ATM phosphorylation at S1981 in the nucleus of individual cells were determined using Cellprofiler software (Carpenter et al., 2006; Kamentsky et al., 2011; McQuin et al., 2018). The NMD efficiency in single cells was calculated by the intensity of EGFP normalized by the intensity of mCherry (EGFP/mCherry ratio). Single dots denote fluorescence intensity of IF and NMD efficiency from single cells. X- or y-axis shows mean intensity of IF or normalized NMD efficiency. No correlation between NMD efficiency and the level of IF detected ATM phosphorylation at S1981 was found in WW expressing cells. The dotted line indicates the regression line. The correlation test between NMD efficiency and the level of IF was performed by Graphpad prism software.

In the NMD fluorescence reporter, the PTC is located downstream of the fluorescence coding sequence in this reporter system, in which the translation of EGFP occurs prior to PTC recognition, the detection of EGFP intensity by flow cytometry did not determine whether the escape resulted from readthrough of the PTC or termination at the PTC with subsequent mRNA degradation. However, the molecular weight of the EGFP-tagged protein and the intensity of band detected by western blotting analysis determined whether there has been readthrough and whether the protein was truncated or full length (**Figure 5**).

**Figure 5.**
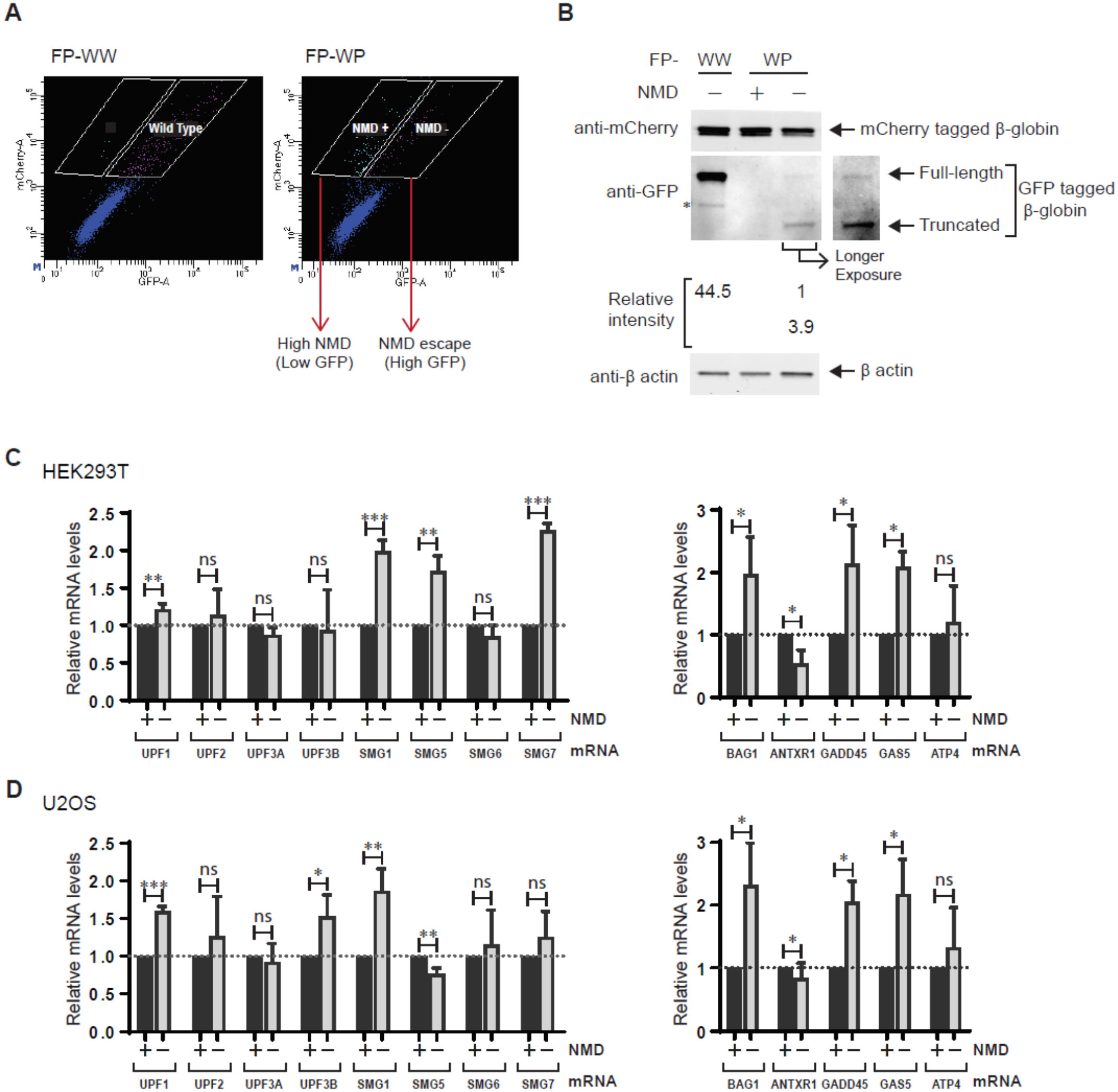
Cell isolation correlated with NMD efficiency. Flow cytometric analysis of FP-WW or FP-WP expressing HEK293T PonA cells. Based on the mCherry and EGFP intensities in FP-WW expressing cell, the cell populations with distinct NMD efficiency (NMD +; higher NMD efficiency, NMD -; NMD escape) were determined and sorted using FACS. White boxes indicate the gate for cell isolation for FACS. Single dots denote fluorescence intensities from single cells. (B) The EGFP and mCherry proteins expressed in isolated cell populations in (A) were detected in western blotting. Same number of cells isolated in (A) were loaded in each lane. EGFP-tagged wild-type β globin was detected in the first lane (FP-WW). In contrast with no detectable bands of EGFP-tagged β globin in second lane (efficient NMD cell population (NMD +) from FP-WP expressing cells), the differential intensities of EGFP bands between lower (truncated EGFP-tagged β globin) and higher (full-length EGFP-tagged β globin) indicated the frequencies of translation termination at PTC and readthrough. Relative intensities of EGFP bands are indicated under the blot detected with anti-GFP antibody. *degradation product. β actin was detected as a loading control. (C) Quantitative mRNA detection of NMD regulators in NMD escape cells. Relative mRNA levels of NMD regulators in the cell population with NMD escape (NMD-) compared to the cell population with efficient NMD (NMD+) in HEK293T ponA (C, left) or U2OS ponA cells (D, left) were determined by RT-qPCR using primers specific for each transcript as indicated above each bar. (D) Quantitative mRNA detection of endogenous NMD-targets (BAG1; BCL2-associated athanogene 1 (Gehring et al. 2005; Wittmann et al. 2006), ANTXR1; ADP Ribosylation Factor Related Protein 1 (Tani et al., (2012)), GADD45; Anthrax toxin receptor 1 (Tani et al., (2012)), GAS5; growth arrest specific 5 (Tani et al., (2013)), ATF4; activating transcription factor-4 (Mendell et al., (2004)) in NMD escape cells. Relative mRNA levels of endogenous NMD targets in the cell population with NMD escape (NMD-) compared to the cell population with efficient NMD (NMD+) in HEK293T ponA (C, right) or U2OS ponA cells (D, right) were determined by RT-qPCR using primers specific for each transcript as indicated above each bar. (C&D) Beta-actin (ACTB) was used as a control. P values were determined using two-tailed unpaired t-tests (***P<0.001, **P<0.01, *P<0.05, ns=not significant). Error bars = Standard deviation from three independent experiments.

The constructs were transiently transfected into HEK293T ponA cells and induced for 24-hours. The specific cell population that escaped from NMD (higher EGFP intensity) or exhibited efficient NMD (lower EGFP intensity) were isolated using FACS (**Figure 5A)**. The protein levels and sizes of EGFP and mCherry in each cell population were determined using western blotting (**Figure 5B)**. Since PTC containing GFP-globin mRNA was degraded efficiently by NMD, a band representing GFP was not found in cells with higher NMD efficiency. In contrast, two distinct molecular sizes of GFP bands were detected in the cells exhibiting NMD escape. The band with higher molecular weight represents full-length GFP-globin protein which resulted from translational readthrough as the identical size of the band was found in the cells expressing EGFP-tagged wild-type Gl. Interestingly the band with smaller molecular weight was detected in the cells showing NMD escape, which represented the truncated EGFP-tagged β-globin proteins resulting from the PTC. The relative intensities of each band indicates that almost 80% of translating ribosomes successfully terminated at the PTC, and 20% engaged in translational readthrough during NMD escape.

In addition to assessing the protein levels of NMD regulators and NMD efficiency (**Figure 3**), we also determined the mRNA levels of NMD regulators by RT-qPCR. The constructs were transiently transfected into HEK293T ponA cells or U2OS ponA cells and the FACS isolated cell populations with the representative NMD efficiencies (as in **Figure 5A**) were subjected to quantitative RT-qPCR analysis. The mRNA levels of selected NMD regulators were correlated with the NMD efficiency. In contrast with the lower protein levels of SMG1 in the cells that have escaped (**Figure 3B**), the relative enrichment of NMD regulators compared with the cells with efficient NMD showed that mRNA levels of UPF1 and SMG1 were higher in both HEK293T ponA and U2OS ponA cells (**Figure. 5C and D left**). The inconsistency between lower protein level (**Figure 3**) and increased mRNA level of SMG1 in the NMD escape cells could be due to the known auto-regulatory feedback of the NMD mechanism: NMD targets mRNAs of its own regulators, most of which contain longer 3’UTRs. Therefore, mRNA of NMD regulators might be stabilized in NMD escape cells. (Huang et al., 2011; Yepiskoposyan et al., 2011). The mRNA levels of UPF2, UPF3A, and SMG6 did not correlate with NMD efficiency in both cell types. The differential correlation of UPF3B and SMG7 mRNA for the two cell types could be explained by cell-type specific auto-regulatory feedback as reported previously (Huang et al., 2011). Moreover, NMD efficiency of endogenous NMD-targets in each FACS-isolated cell population were assessed as a control since NMD escape may be transcript-dependent (or reporter-dependent). Using RT-qPCR analysis, we detected higher mRNA levels of endogenous NMD targets, except ANTXR1 mRNA, in both cell types that escape compared to the cells with efficient NMD. (**Figure 5C and D Right**). This suggests that NMD escape has a general effect on the transcriptome.

## Discussion

In this study, we established single-cell detection of NMD efficiency using a NMD reporter construct containing two ORFs, which express wild-type and PTC containing mRNA bi-directionally in the same cells. Simultaneous detection of wild-type and PTC-containing mRNA in single cells minimizes the internal variables related to gene expression other than NMD, thus it provides the precise detection of NMD efficiency in every cell. Using this reporter system, single mRNA detection with smFISH revealed a variable range of NMD efficiencies in the cell population, providing evidence of cell-to-cell heterogeneity of NMD. This simultaneous detection of two genes, wild-type and mutant, provides a general application for single-cell biology. The intercellular variability in cell populations has been acknowledged by advanced technology such as single-RNA seq, however general biochemical assays addressing post-transcriptional regulation are not capable of single-cell detection. In this system, the expression of wild-type mRNA in the same cells verifies the proper biological activity representing an internal control in individual cells. Together with the single-molecule mRNA imaging tools such as smFISH and live imaging using the MS2 system, the direct comparison of wild-type and mutant mRNAs in same cells can address multiple points of regulation (e.g. transcription, abundance, mRNA localization). A dual fluorescence reporter is compatible with the flow cytometric detection and cell isolation using FACS, enabling the investigation of rare but informative cells that are not detectable in the general biochemical approach.

NMD escape is a general biological characteristic for most NMD targets. Previous studies have also supported that conditional NMD inhibition is a part of rapid post-transcriptional regulation in response to several environmental stimuli or physiological changes. Global translational inhibition is known to be a mechanism of NMD inhibition under several cellular stress conditions (Brown et al., 2011; Brumbaugh et al., 2004; Masse et al., 2008; Nergadze et al., 2004; Oliveira et al., 2008). Therefore, we characterized NMD escape while cells were undergoing active translation, using a fluorescence protein indicator.

Our dual-fluorescence NMD reporter system provides a conventional single-cell detection of NMD. The detection of NMD efficiency using this system can be accomplished by simple fluorescence detection without requiring RNA purification or fluorescence labelling. In this study, we correlated NMD efficiency with NMD regulators to identify the factors associated with variability. Using a fluorescence reporter for detecting NMD and IF for specific NMD regulators, we identified that SMG1 is correlated with NMD variability. The correlation of NMD efficiency with ATM phosphorylation, known to be a partner of SMG1 during DSB repair, indicated a possible a link between DNA damage response and NMD variability. Although pharmacological treatment inducing DNA damage inhibits NMD (Yamashita et al., 2001), a causal relationship between DNA damage with reduction of SMG1 protein levels in cells is still unclear since downregulation of SMG1 activates p53 phosphorylation and mediates DNA damage response without inducing genotoxic stress (Brumbaugh et al., 2004).

One of the advantages of the dual-fluorescence NMD reporter system is its compatibility in the isolation of differential NMD efficiencies by FACS, which permits further investigation into a minor cell population. Since PTC recognition is a key event to trigger efficient NMD, we investigated whether cells escape from NMD with PTC recognition or readthrough. Isolation of NMD escape cells followed by western blotting determined that 20% of translating ribosomes readthrough the PTC in the cell population that escapes surveillance. This approach provides a sensitive assessment to elucidate the pathways whereby the PTC was recognized (or not) and decay was not triggered. In addition, the approach described here can potentially expand into future studies that investigate the role of NMD in many biological phenomena. In fact, NMD plays a significant role in cell differentiation, memory consolidation, virus infection, tumorigenesis, and immune response (Lykke-Andersen and Jensen, 2015; Ottens and Gehring, 2016).

Lastly, a study regarding NMD escape potentially provides an approach for drug discovery targeting of nonsense suppression. Inducing translational readthrough at the PTC, an attractive strategy for nonsense-associated disease-causing genes. The major limitation of nonsense suppression therapy is a significant reduction of nonsense transcripts by NMD. Our approach may offer a sensitive assay for translational readthrough that will provide valuable insights into unknown regulatory pathways resulting in NMD escape.

## Author Contributions

Conceptualization, H.S.; Methodology, H.S; Formal Analysis, H.S; Investigation, H.S., R.H.S.; Resources, R.H.S.; Data Curation, H.S.; Writing - Original Draft, H.S. Writing - Review & Editing, R.H.S. Visualization, H.S. Supervision, R.H.S. Funding Acquisition R.H.S.

## Acknowledgments

This work was supported by NIH grant R01 NS083085 (R.H.S). We thank members of the Singer laboratories for discussions, the Einstein FACS and Genomics cores. We also thank L. Maquat for anti-UPF1 antibody. The authors declare no competing financial interests.

