## Supplementary materials for "Cellular variability of nonsense-mediated mRNA decay"

**This PDF includes:**

Supplementary Figures S1 to S6
Supplementary Table S1 to S2
Methods

### Supplemental Results

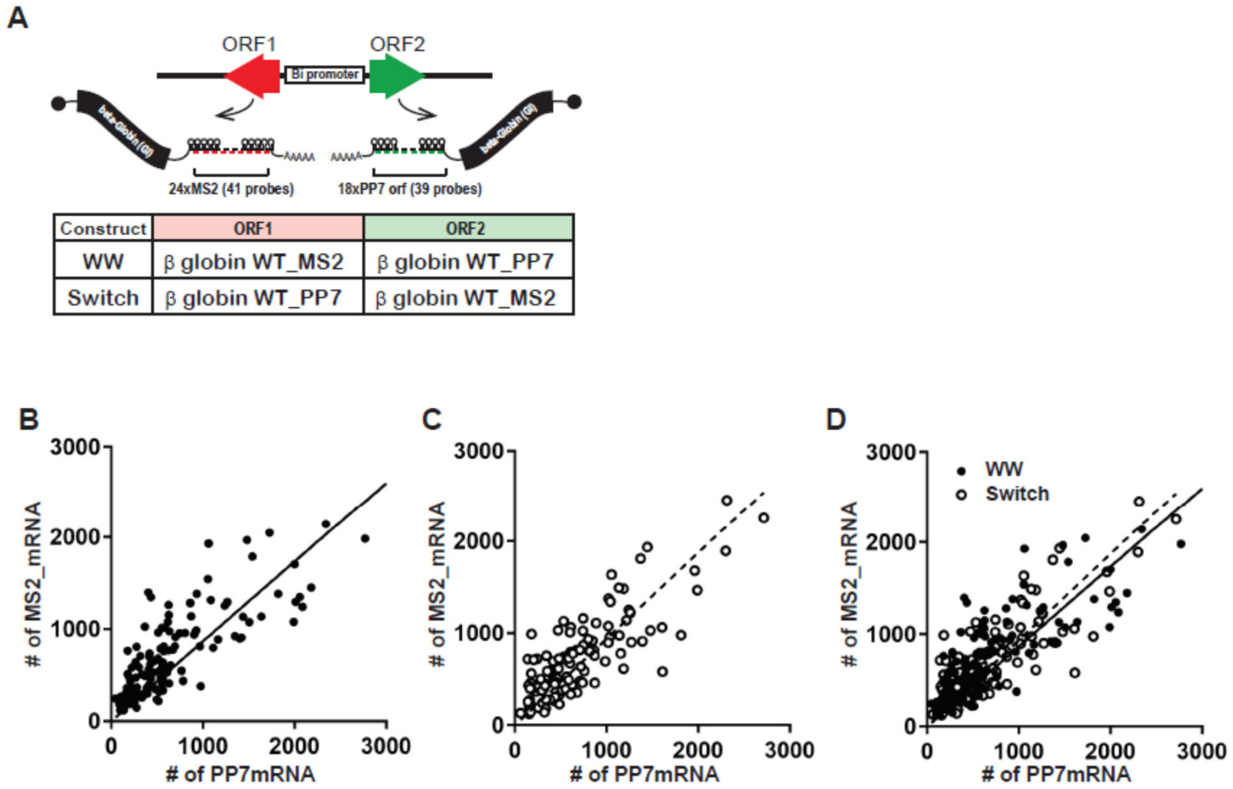


**Supplemental Figure S1.** Comparable expression of β-globin mRNA with MS2 or PP7 under bidirectional promoter. (A, upper) Schematic of the smFISH to detect β-globin (Gl) mRNA expressing under PonA bidirectional promoter. These transcripts contain MS2 or PP7 sequences in the 3'UTR that hybridize with unique FISH probes. (A, lower) The table shows the constructs that we used in this experiment. (B-D) Linear correlation of Gl_MS2 and Gl_PP7 gene expression under bidirectional promoter and switching ORF does not affect their expressions. Lines or dotted lines indicate the regression lines of WW or Switch expressing cells. Total number of Gl_MS2 and Gl_PP7 mRNAs in steady-state U2OS PonA was detected using smFISH and mRNA spots were counted in cells expressing WW (B&D) or Switch (C&D) construct[26]. Single dots denote the number of MS2 (*y*-axis) and PP7 (*x*-axis) mRNAs from single cells. $n_{cells}$ =116 and 113 for WW and Switch constructs. Two data sets from U2OS PonA cells expressing WW or Switch were superimposed in (D). Pearson r=0.81 and 0.75 for WW and Switch constructs.

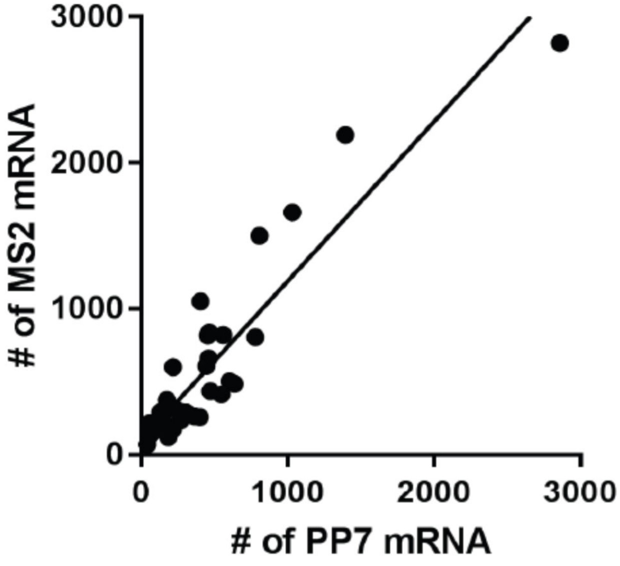


**Supplemental Figure S2.** Linear correlation of GI_MS2 and GI_PP7 gene expression under bidirectional promoter after 1hr PonA induction. The total number of GI_MS2 and GI_PP7 mRNAs were detected using smFISH and single mRNA spots were counted in U2OS PonA cells expressing WW. Single dots in scatter plot denote the number of MS2 (y-axis) and PP7 (x-axis) mRNAs from single cells. $n_{cells}$ =35. Pearson r=0.9175, R squared= 0.84. Peason r and R square were calculated by Graph Pad Prism software. Lines indicate the regression lines of WW.

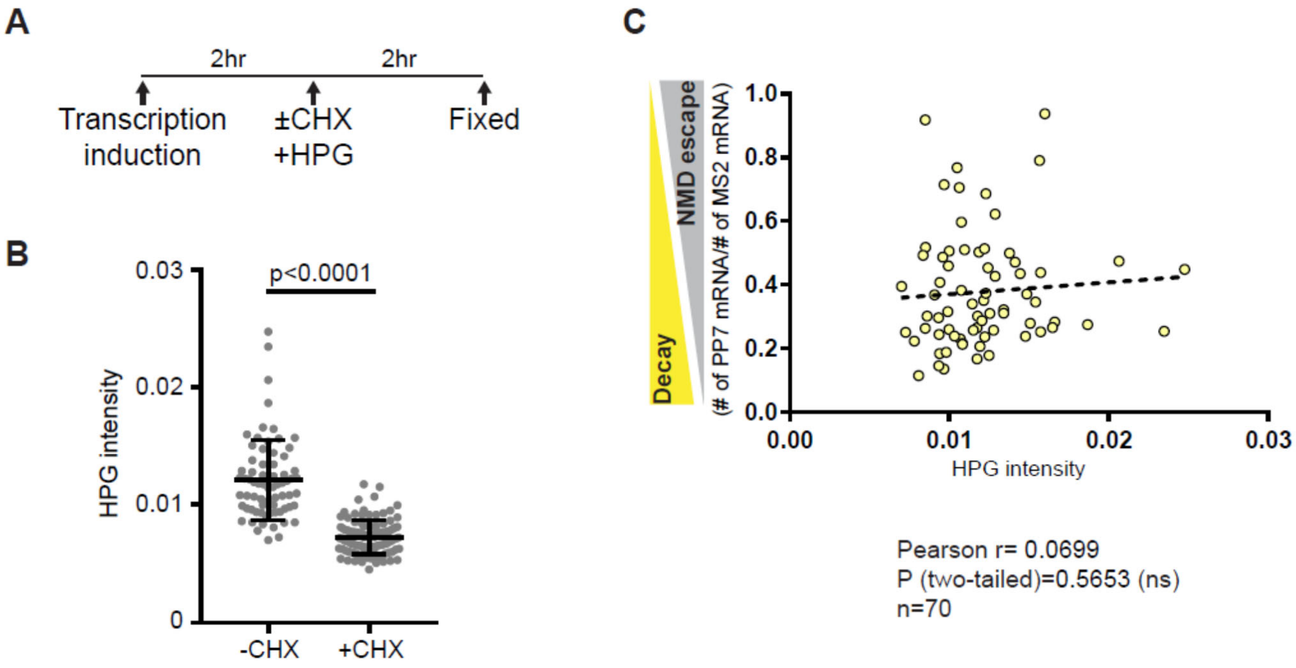


**Figure S3. NMD efficiencies do not correlate with translation activity.**
Single-cell detection to correlate translation activity and NMD in single cells. Translation activity in each cell was determined using the Click-it HPG system. (A) The timeline of transcription induction by supplement of PonA and labeling of nascent peptides with HPG incorporation. CHX, Cycloheximide; HPG, L-Homopropargylglycine (B) Click-it HPG successfully detected translation reduction. The translation inhibition using CHX decreased HPG incorporation detected by the Click-it labeling system. (C) No correlation was found between NMD escape and translation activity. The level of NMD escape in single cells was calculated by the number of PTC-containing GI mRNAs normalized by the number of wild-type GI mRNAs that were detected with smFISH. The dotted line indicates the regression line. The correlation test between NMD efficiency and the level of HPG was performed by Graphpad prism software.

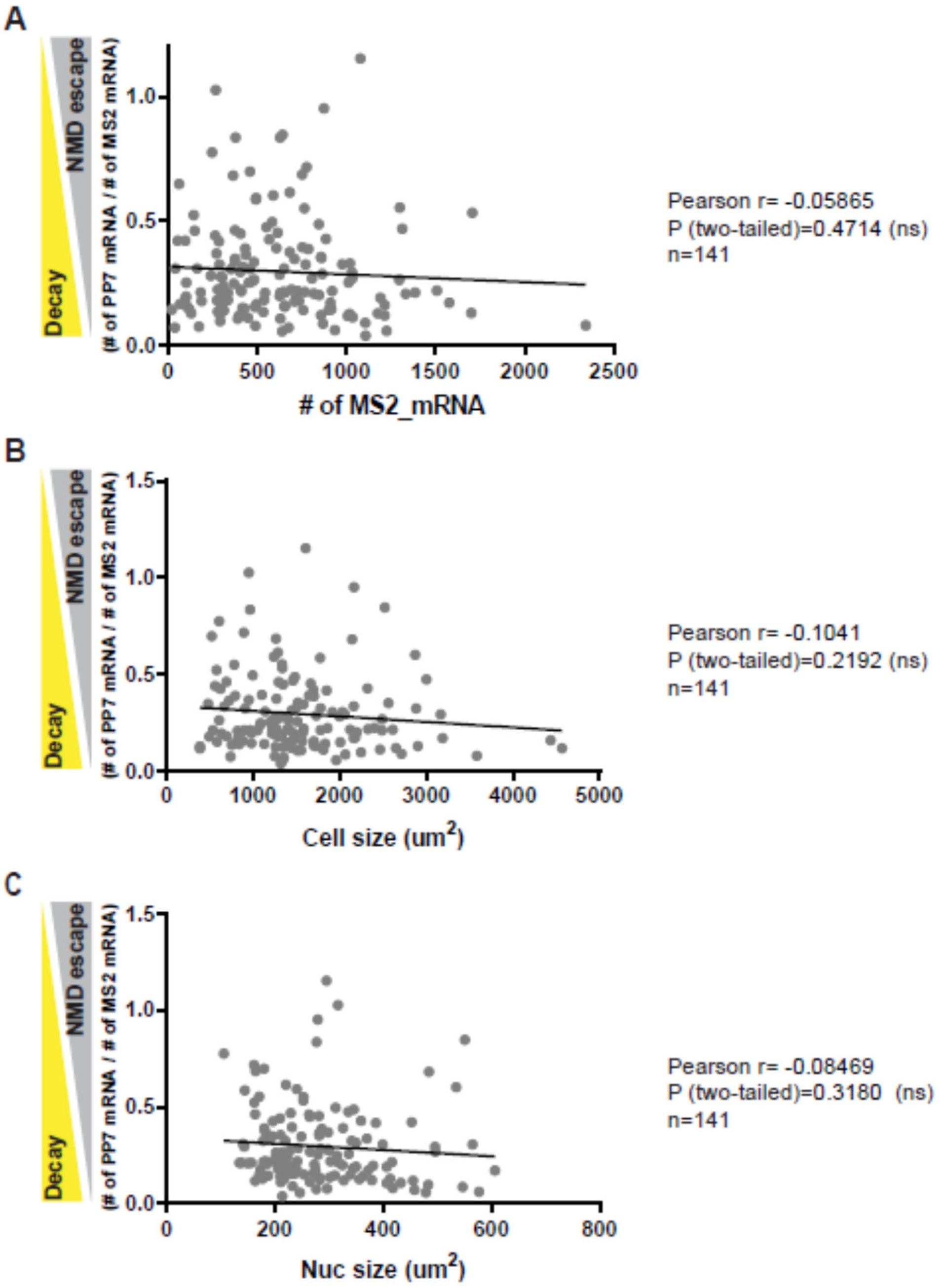


**Supplemental Figure S4.** The level of NMD escape does not correlate with the level of mRNA expression, the cell size, or nuclear size. The level of NMD escape in single cells was calculated by the number of PTC-containing GI PP7 mRNAs normalized by the number of wild-type GI MS2 mRNAs. (A) The level of mRNA expression was estimated by the level of GI WT MS2 mRNA which is not degraded by NMD. (B) The cell size was measured using FISH-quant based on the cell outline area. (C) The area of the nucleus was defined using the threshold of DAPI staining. Correlation test between NMD efficiency and (A) the level of mRNA expression (Pearson r=-0.05865), (B) cell size (Pearson r=-0.1041) or (C) nuclear size (Pearson r=-0.08469) was performed by Graphpad prism software.

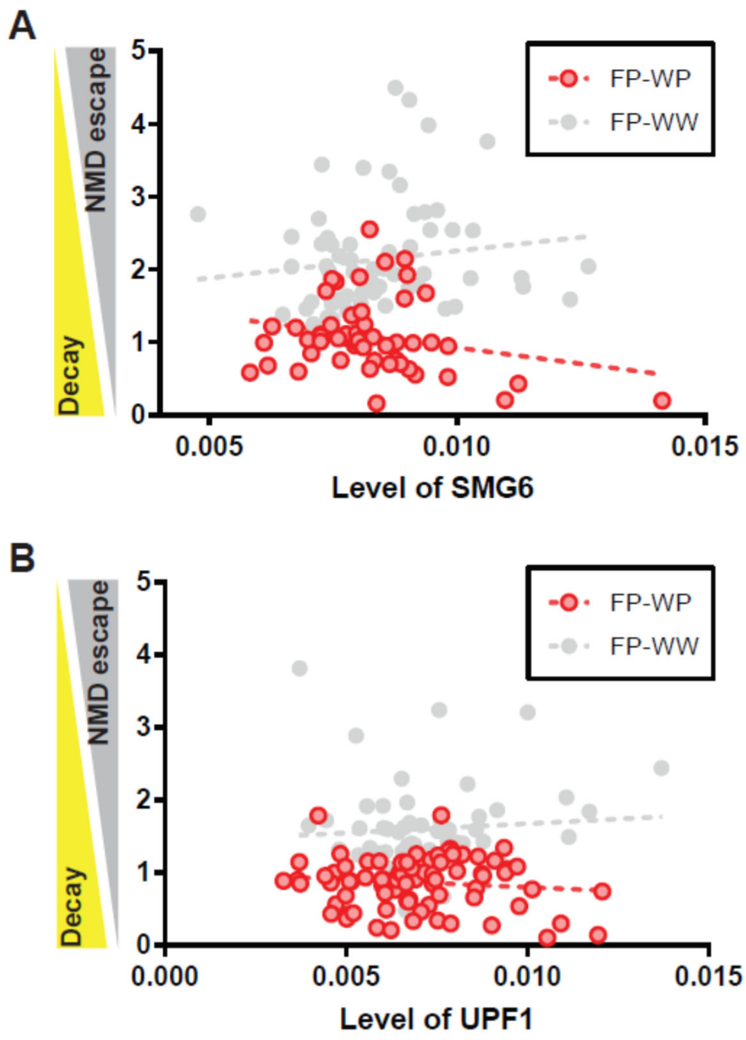


Pearson r= -0.2396
P (two-tailed)=0.0781 (ns)
n=55

Pearson r= -0.1115
P (two-tailed)=0.3001 (ns)
n=83

**Supplemental Figure S5.** The level of NMD escape does not correlate with the protein level of (A) SMG6 and (B) UPF1. The NMD efficiency in single cells was calculated by the intensity of EGFP normalized by the intensity of mCherry (EGFP/mCherry ratio). Single dots denote fluorescence intensity of IF and NMD efficiency from single cells. X- or y- axis shows mean intensity of IF or normalized NMD efficiency. No correlation between NMD efficiency and the level of IF detected ATM was found in WW expressing cells. The dotted line indicates the regression line. The correlation test between NMD efficiency and the level of IF was performed by Graphpad prism software.

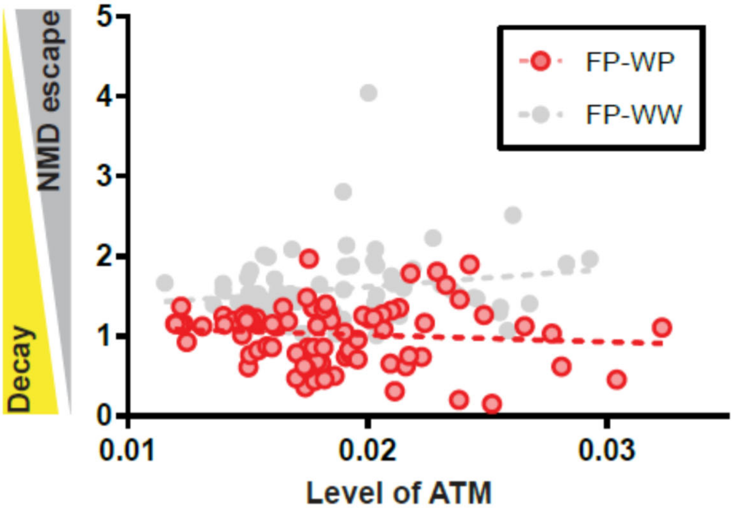


**Supplemental Figure S6.** No correlation of NMD efficiency with the level of ATM.
The level of ATM was determined by immunofluorescence using anti-ATM antibody followed by Alexa Fluor 647 linked anti-rabbit IgG. The intensity of mCherry, EGFP, DAPI-stained nuclei, or the level of ATM was detected using fluorescence microscopy. Bar = 10 µm. No correlation was found between NMD efficiency and the level of ATM. Pearson r=-0.09584, P (two-tailed) =0.3947, n=81. Mean intensity of EGFP, mCherry and IF representing the level of ATM in the nucleus were determined using Cellprofiler software [31-33]. The NMD efficiency in single cells was calculated by the intensity of EGFP normalized by the intensity of mCherry (EGFP/mCherry ratio). Single dots denote fluorescence intensity of IF and NMD efficiency from single cells. X- or y- axis shows mean intensity of IF or normalized NMD efficiency. No correlation between NMD efficiency and the level of IF detected ATM was found in WW expressing cells. The dotted line indicates the regression line. The correlation test between NMD efficiency and the level of IF was performed by Graphpad prism software.

### Supplementary Table S1: Probe list for smFISH

| MS2 Probe Sequence (5' to 3') |
|---|
| tgattgtgaagtgtcgggtg |
| tccacccttgtgtattgtac |
| tgtaatgtgtctggagggtg |
| gcttctgtttgattggattt |
| gatggtgattccttgttgta |
| gtatattgcacagggaatcc |
| gatattcgggaggcgtgatc |
| acgcactgaattcgaaagcc |
| attcgactctgattggctgc |
| ctcttcgcgaaagtcgactt |
| taagaatggcgcgaaggctg |
| gtaggggagagtgtggtttg |
| caggaacgctgatgctgttc |
| ttttcttgagttgggtactg |
| tgatgctgcatggggacata |
| ttggggatgtattcttgggg |
| ttggtgctcggatgtgattt |
| aagaaacaacactccgagcc |
| atggagggtttgtccagttg |
| tttgtcttgttggtgagagt |
| ctgatgctgcttcgagaaga |
| gtatgctcgagtgtttcgaa |
| gatcgtccacccaagaaata |
| aattcgtgagagcatgggtg |
| tcgtattggacgtggaacga |
| tcgtgatcccgaaaggtaag |
| atcgtgcatgcttgaatgtc |
| gttgagacttgtggagcatg |
| tgaacccatttggtagtttc |
| tttgaggtaggagtgggttc |
| ttgccagttttgtgggaaga |
| tttggtatgttggaatgggc |
| gatgctgtaccagtaattgt |
| tagtagtgagagatgtgggc |
| tgctgaacggtttggttttt |
| ttgatttttccgtgtgtacc |
| gtctttcgtatttgtaaacc |
| ttgcgctggacgaaagcgtg |
| ccgtcggatgtttttcgtaa |
| ggttgtaagtttgtgggttg |

| PP7 Probe Sequence (5' to 3') |
|---|
| tacaaaattggctctcggcg |
| ggttttgtcgagaaactggc |
| tgtatacatagcggagggac |
| tgaccttcgatgactgtctg |
| catatggtctgctcccatac |
| taatagccgcgacattcttt |
| cctctgtccgtgtacaaaaa |
| acaggtacagacggacgagg |
| agtaataagcgctgagcgcg |
| atatcctctgtcggcgtatc |
| gacatttgggaacagtcgga |
| tataaggacggaggccatac |
| agtaaaggttcttggtcgac |
| tctctgggaccagatacaag |
| acgcgttcccaaatcttttg |
| ttgtatgattggcctcggag |
| tgcgcacagagcataatact |
| ttgtacataaccggagcctc |
| tcgcgggtacacaaactgat |
| catacggtgtgctggtaagt |
| tatatggacagggacaggga |
| ggtctgtcgggatacataac |
| ctcgacaaataccgacgacc |
| acaattttgtacttgcgtgt |
| catatgtcctgcgagatatc |
| aatagagatctgctctcgcg |
| atatcctctggtgccgatag |
| aagatgctgaggcaccgagg |
| ttggacagtgacagcacatc |
| atatgctctgctggcttgaa |
| tgtaatgacagtggagccag |
| gattggagaagatcaggtcc |
| atgtagtcggagggagcgag |
| tagtctctcagcacttcgag |
| catatcgctgaccctttgat |
| gaacgagtagtaggggtcgc |
| cgactctgcgttgaagagta |
| tttgtatctctgatggacgc |

| |
|---|
| ctgaggtgtttgatgtacgg |

**Supplementary Table S2: Primer list for RT-qPCR**

| | Forward | Reverse |
|---|---|---|
| UPF1 | AGATCACGGCACAGCAGAT | TGGCAGAAGGGTTTTCCTT |
| UPF2 | TCTCACCTGAGGACCAGTGTAC | AGCTGGAGGTGGGTTGCAGTAG |
| UPF3A | TACTGGAGGTGGCAAGCAGGAA | CCTGTGCTCTTTATCACTGCCG |
| UPF3B | GAAAGAGCCAGTGGGCAAAGTTG | CGAAGTATGCGCTCCTGATCTC |
| SMG1 | TCGAAGTCAAGAACACGTTGA | GGGTGATGCAAAACTCACTAAA |
| SMG5 | CACTAAGCGGCCGCTACTGAC | TCTATGCGGCCGCCTAACGTCTTGGCAACAAAGGGAC |
| SMG6 | GATGGTCTTGCCATTCGCAGCA | TCGCTGTATCACTGGCTTGCTC |
| SMG7 | TTTCAGGAGGCAGTGGTGGATG | CAAACTCCTCTGGAAGTGGTGTC |
| BAG-1 | GTGAACCAGTTGTCCAAGACCTG | CAAGTGCTGACAACGGTGTTTCC |
| ANTXR1 | ATGCCTTGTGGGTCCTACTG | GAGGTGTGGTAGGCGTTGTT |
| GADD45A | GGAGGAATTCTCGGCTGGAG | CGTTATCGGGGTCGACGTT |
| ATF4 | TCAACATCGTGCGGGTGTCG | CCCGGCTTTCTTCGCAGTAG |
| GAS5 | CTTGCCTGGACCAGCTTAAT | CAAGCCGACTCTCCATACCT |
| ACTB | TCCCTGGAGAAGAGCTACG | GTAGTTTCGTGGATGCCACA |

### Methods

#### Cell Lines and Tissue Culture

Human U2OS PonA cell line was generated by stable transfection of pERV3, which expresses synthetic VP16-glucocorticoid/ecdysone receptor (VgEcR) and retinoid-X-receptor (RXR) that are required for induction for the transcription of PonA promoter as described previous study[22]. Human embryonic kidney 293T PonA cell was generated by stable transfection of pERV3-zeocin vector, which is replaced neomycin drug selection gene in pERV3 to Zeocin gene due to the neomycin resistance of HEK293T cells. Both cell lines were grown at 37°C and 5% $CO_2$ in Dulbecco's Modified Eagle's Medium (DMEM) supplemented with 4.5 g/l of glucose, 10% FBS and 1% penicillin-streptomycin.

#### Cell transfections, single molecule fluorescence in situ hybridization (smFISH), immunofluorescence (IF) and fixed cell image acquisition

U2OS PonA cells were transiently transfected with Bi-directional PonA construct, pFRT-PonA-BI-GI-WT-24xMS2-WT-18xPP7 (WW), pFRT-PonA-BI-GI-WT-24xMS2-PTC-18xPP7 (WP), or PonA-BI-GI-WT-18xPP7-WT-24xMS2 (Switch) with the Amaxa Nucleofector system, then spread onto collagen-coated coverslips. One day after transfection, transcription was induced by supplement with 20nM PonA for 24-hours (steady-state) in Figure 1 and S1, and 1-hr for Supplemental Figure S2. The cells were fixed in 4% paraformaldehyde for 15 minutes at room temperature, then permeabilized with 0.5% Tween in PBS for 15 minutes at room temperature. smFISH was performed as previously described[24]. In Figure 2-5, U2OS or HEK293T PonA cells were transiently transfected with pFRT-PonA-BI-GI-GFP-WT-mCherry-WT (5FP-WW) or pFRT-PonA-BI-GI-GFP-WT-mCherry-PTC (5FP-WP). Transcription was induced by supplement with 20nM PonA for 24-hours (steady-state). The cells were fixed in 4% paraformaldehyde for 15 minutes at room temperature, then permeabilized with 0.5% Tween in PBS containing 3% BSA for 15 minutes at room temperature. The cells were incubated with anti-UPF1 (1/1000 dilution, gift from Dr. Maquat), anti-phospho UPF1 (1/200 dilution, Millipore, 07-1016), anti-SMG1 (1/200 dilution, Cell Signaling 9149S) or anti-SMG6 (1/200 dilution, invitrogen PA5-60165), anti-ATM (1/200 dilution, Novus Biologicals, NB100-104SS), anti-ATM S1981P (1/500 dilution, Active Motif #39530) for 1-hour at room temperature. After 10-min washing with PBS for three time at room temperature, the cells were incubated with Alexa Fluor 647 secondary anti-rabbit IgG antibody (Thermo Fisher Scientific, car# A21245), or anti-mouse IgG antibody (Thermo Fisher Scientific, car# A21236) at a concentration of 2 μg/mL in PBS for 1-hour. After 10-min washing with PBS for three time at room temperature, the cells were mounted with Prolong Diamond Antifade Mountant with DAPI (Thermo Fisher Scientific. P36966). Images were acquired with an Olympus BX61 widefield, epifluorescent microscope using a 60X 1.4 PlanApo objective. Filter sets were used for DAPI (Semrock model DAPI-5060C-Zero), Cy3 (Chroma model 41007), Cy3.5 (Chroma model SP103v1), and Cy5 (Semrock model Cy5-4040C-Zero), with an EXFO X-Cite Series 120 PC metal halide light source, Photometrics Cool SNAP HQ CCD camera, Olympus Type-F immersion oil (nd 1.516) and Molecular Devices Metamorph acquisition software. Cells were optically sectioned using a 0.25 μm Z step, spanning a 10.0 μm Z depth in total. Exposure times of 20 to 500 ms were typically used to acquire each plane in the Cy3, Cy3.5 and Cy5 channels, and ~12 ms were used to acquire each plane in the DAPI channel. All the probes used in smFISH in this study are provided in Supplementary Table 1.

#### Detection of translational activity using HPG labelling

L-Homopropargylglycine (HPG) labelling was performed as describe in manufacturer's instructions. Briefly, U2OS PonA cells were transiently transfected with Bi-directional PonA construct PonA-BI-GI WT-24xMS2 PTC-18xPP7 (WP) with the Amaxa Nucleofector system, then spread onto collagen-coated coverslips. One day after transfection, 25 μM HPG (from Click Chemistry Tools, Product No.1067) was added in the methionine-free DMEM medium (Life Technologies Corporation, Part Number 2101302) with or without cycloheximide (50 μg/ml) treatment for 2 hours after transcription induction by supplement with 20nM PonA for 2 hours. The cells were fixed in 4% paraformaldehyde for 15 minutes at room temperature, then permeabilized with 0.5% Tween in PBS for 15 minutes at room temperature. HPG was fluorescently labeled using click-it technology (Click-&-Go Plus

488 Labeling Kit, Product No.1314 from Click Chemistry Tools), then smFISH was performed as previously described[24].

### RT-qPCR

Total cell RNA extraction, DNase I treatment, and reverse transcription were performed as described previously (Sato and Maquat 2009). Amplifications were performed using Power SYBR® Green PCR Master mix (Applied biosystems, REF 4367659). Primers used in this study are listed in Supplementary Table 2. All samples were analyzed in three times, and the levels of mRNAs were normalized to the level of β-actin mRNA. Relative mRNA levels were determined from CT values according to the ΔΔCT method (Applied Biosystems).

### Flow cytometry analysis and cell sorting

Flow cytometry detection was performed using cell analyzer LSRII (BD Biosciences) and fcs files were analyzed by FlowJo (BD Biosciences). In Figure 5, specific cell populations were gated and isolated using FACS Aria II (BD Biosciences).

### Western blotting

The cells isolated by FACS were lysed using hypotonic lysis buffer (Lykke-Andersen et al., 2001) and cell lysate was electrophoresed using NuPAGE system (4%–12%, Thermo Fisher Scientific), then transferred to nitrocellulose membrane using iBlot 2 (Thermo Fisher Scientific). The membrane was probed with anti-mCherry (Novus Biologicals, NBP2-25157SS), anti-GFP (Roche, REF 11814460001), or anti beta-actin (Sigma, A1978) antibody following anti-rabbit IRDye® Secondary Antibodies. The signal was detected using Odyssey imaging system (LI-COR Biosciences).

### Data analysis

The number of mRNA spots were detected and counted in the smFISH images using FISH-quant[26]. Standard deviation, two-tailed paired t-tests, pearson and R squared were calculated with GraphPad Prism. The levels of HPG and NMD regulators labelled with IF, which was detected fluorescent microscope, were measured using CellProfiler based on the area of individual cells determined by CellProfiler.
